# YAP–TEAD-driven CPA4 promotes NF2-deficient meningioma growth

**DOI:** 10.64898/2026.08.05.743013

**Authors:** Kentaro Mineji, Katerina Petrosky, Ryosuke Otsuji, Yasuhide Makino, Yuji Kibe, Eita Uchida, Daichi Hagita, Neha Singaravelan, Yukitomo Ishi, Shigeru Yamaguchi, Long- Sheng Chang, Samantha Gadd, Rintaro Hashizume

**Affiliations:** Department of Pediatrics, University of Alabama at Birmingham, Birmingham, AL, USA; Division of Pediatric Hematology and Oncology, Children’s of Alabama, Birmingham, AL, USA; Department of Neurosurgery, Faculty of Medicine and Graduate School of Medicine, Hokkaido University, Sapporo, Japan; Center for Childhood Cancer, Abigail Wexner Research Institute at Nationwide Children’s Hospital and Department of Pediatrics, The Ohio State University, Columbus, OH, USA; Quantitative Science Pillar, Stanley Manne Children’s Research Institute, Ann & Robert H. Lurie Children’s Hospital of Chicago, Chicago, IL, USA; O’Neal Comprehensive Cancer Center, University of Alabama at Birmingham, Birmingham, AL, USA

## Abstract

Neurofibromin 2 (NF2) deficiency is a driver of meningioma and other cancers, yet transcriptional effectors that sustain NF2-deficient tumors remain poorly defined. We identify carboxypeptidase A4 (CPA4) as an effector of YAP–TEAD signaling in NF2-deficient meningioma. Transcriptomic profiling identified CPA4 as a consistently upregulated effector. Across patient cohorts and specimens, CPA4 expression was enriched in *NF2*-mutant and chromosome 22q-deleted meningiomas and associated with higher tumor grade and chromosome 1p loss. CPA4 depletion impaired proliferation, disrupted cell-cycle, DNA-replication, and DNA-repair programs, suppressed intracranial tumor growth, and prolonged survival. Integrated epigenomic and functional assays identified CPA4 as a direct YAP–TEAD transcriptional target. CPA4-high meningioma models exhibited preferential sensitivity to YAP–TEAD inhibition, while verteporfin and the clinical-stage TEAD inhibitor VT3989 reduced CPA4 expression, suppressed orthotopic tumor growth, and prolonged survival. These findings uncover a targetable YAP–TEAD–CPA4 dependency in NF2-deficient meningioma and identify CPA4 as a potential biomarker for TEAD-directed therapy.

**STATEMENT OF SIGNIFICANCE:** CPA4 links NF2 loss to oncogenic YAP–TEAD transcription, sustains meningioma growth, and marks tumor sensitivity to pharmacologic TEAD inhibition. These findings establish CPA4 as a tumor-promoting effector and potential biomarker of an actionable pathway shared across NF2- deficient cancers.

## INTRODUCTION

Meningiomas are the most common primary intracranial tumors (1). High-grade meningiomas account for approximately 20% of cases and carry substantial risks of recurrence and disease-specific mortality (2). Neurofibromin 2 (NF2) is the most frequently altered tumor suppressor gene in meningioma, with genetic alterations detected in approximately 40–50% of tumors (3). In addition to genetic mutations, epigenetic silencing, including aberrant methylation of the *NF2* promoter, may further contribute to functional NF2 loss (4). NF2-deficient meningiomas exhibit genomic instability and are associated with higher tumor grade, increased recurrence, and poor clinical outcomes (5, 6). DNA methylation profiling has also identified an unfavorable meningioma subgroup characterized by low *NF2* expression (7). Collectively, these findings establish NF2 deficiency as a major driver of meningioma and underscore the need to define the downstream dependencies that sustain these tumors (8).

*NF2* encodes merlin, an upstream suppressor of Hippo signaling. Loss of *NF2* disrupts Hippo pathway control, enabling nuclear YAP/TAZ accumulation and TEAD-dependent transcription that promotes tumor growth (9–12). This signaling state is a defining feature of NF2-deficient cancers and has created strong interest in direct TEAD inhibition. VT3989, a first-in-class oral TEAD palmitoylation inhibitor, has provided early clinical proof of concept for pharmacologically targeting YAP–TEAD signaling in a phase I/II study of advanced solid tumors, particularly mesothelioma (9). However, the tumor-promoting transcriptional effectors that connect NF2 loss to YAP–TEAD activity remain incompletely defined, and validated biomarkers of pathway activity and response to TEAD inhibition are lacking.

Through comparative transcriptomic profiling of NF2-deficient meningioma models, we identified carboxypeptidase A4 (CPA4) as a consistently upregulated effector of NF2 deficiency. CPA4 is a metallocarboxypeptidase implicated in tumor progression and poor outcome in several cancers (13–16), but its function in meningioma and its relationship to NF2–YAP–TEAD signaling have not been established. These observations raised the possibility that CPA4 represents a disease-relevant transcriptional effector of oncogenic YAP–TEAD signaling and a marker of therapeutic vulnerability.

Here, we show that CPA4 expression is enriched in *NF2*-mutant and chromosome 22q-deleted meningiomas and is associated with higher tumor grade and chromosome 1p loss. Inducible CPA4 depletion impaired proliferation, disrupted cell-cycle, DNA-replication, and DNA-repair programs, suppressed intracranial tumor growth, and prolonged survival. Integrated epigenomic and functional analyses established *CPA4* as a direct transcriptional target of YAP–TEAD. CPA4- high meningioma models were preferentially sensitive to YAP–TEAD inhibition, and treatment with verteporfin or VT3989 reduced CPA4 expression, suppressed orthotopic tumor growth, and prolonged survival (9, 17, 18). Together, these findings define a previously unrecognized, targetable YAP–TEAD–CPA4 dependency in NF2-deficient meningioma and identify CPA4 as a potential biomarker for TEAD-targeted therapy.

## RESULTS

### NF2 restoration identifies CPA4 as a candidate downstream effector in NF2-deficient meningioma

To identify transcriptional effectors sustained by NF2 loss, we introduced an NF2 open reading frame (ORF) into the NF2-deficient meningioma cell lines CH157 and KT21-MG1-Luc5 (hereafter KT21-MG1) and profiled the resulting transcriptional changes by RNA sequencing (Figure 1). Integrated analysis identified 234 shared downregulated and 150 shared upregulated genes following NF2 restoration (Figure 1A). Because effectors sustained by NF2 deficiency should decrease after restoration, we prioritized the shared downregulated genes. Heatmap analysis demonstrated consistent transcriptional changes following NF2 restoration in both models and identified *CPA4* among the genes suppressed in *NF2* ORF-expressing cells (Figure 1B). Volcano plots further showed marked *CPA4* downregulation in both cell lines (Figure 1C), and box-plot analysis confirmed a significant reduction in *CPA4* expression following NF2 restoration in each model (Figure 1D). *CPA4* also ranked among the ten most strongly downregulated transcripts in each cell line (Supplementary Table S1). Based on this consistent regulation and its reported tumor-promoting roles in other cancers (13–15), we further investigated CPA4 as a candidate downstream effector of NF2 deficiency.

**Figure 1.**
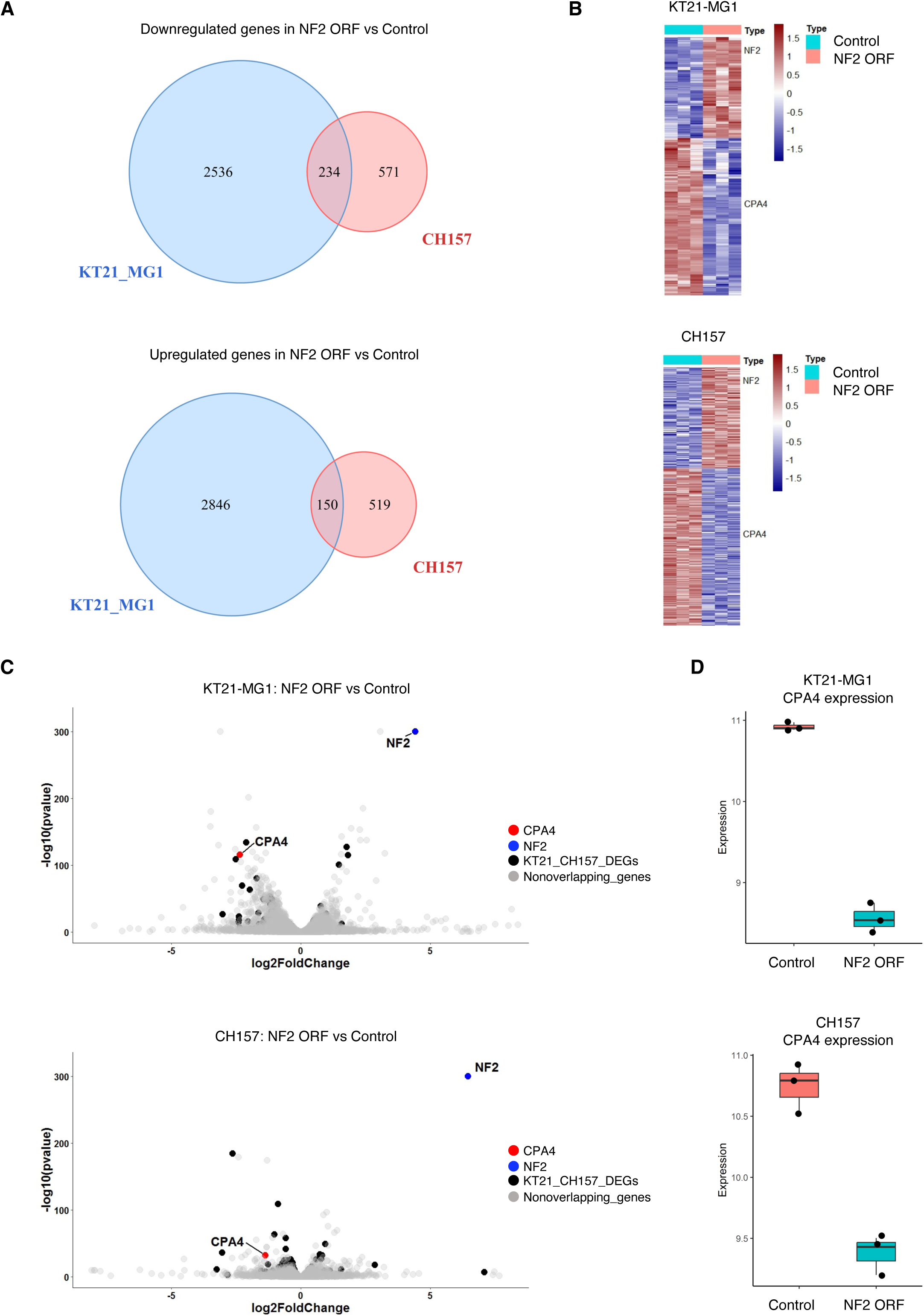
NF2 restoration identifies CPA4 as a candidate downstream effector in NF2-deficient meningioma. (A) Venn diagrams showing shared downregulated and upregulated genes after NF2 ORF expression in CH157 and KT21-MG1 cells (adjusted *P* < 0.05). (B) Heatmaps of differentially expressed genes after NF2 restoration (three biological replicates per condition). (C) Volcano plots showing differential expression after NF2 restoration. CPA4, red; NF2, blue; genes differentially expressed in both models, black; nonoverlapping genes, gray. (D) CPA4 expression after NF2 restoration. Box plots show normalized expression.

### NF2 restoration suppresses CPA4 expression and NF2-deficient meningioma growth

We next examined whether NF2 restoration suppressed CPA4 protein expression and tumor growth. Immunoblotting confirmed robust NF2 protein expression and markedly reduced CPA4 protein abundance in both CH157 and KT21-MG1 cells transduced with lentiviruses expressing an *NF2* ORF (Figure 2A). NF2 restoration also significantly reduced cell proliferation and clonogenic growth in both models (Figure 2B, C). We then orthotopically implanted control and *NF2* ORF-expressing CH157 cells in mice. NF2 restoration suppressed intracranial tumor growth, as assessed by bioluminescence imaging (BLI), and prolonged survival (Figure 2D, E). Tumor immunohistochemistry (IHC) showed a significant reduction in Ki-67-positive cells (Figure 2F) and confirmed increased NF2 expression and reduced CPA4 expression (Figure 2G). Together, these findings link restoration of NF2 function to CPA4 suppression and growth inhibition across cellular and orthotopic models.

**Figure 2.**
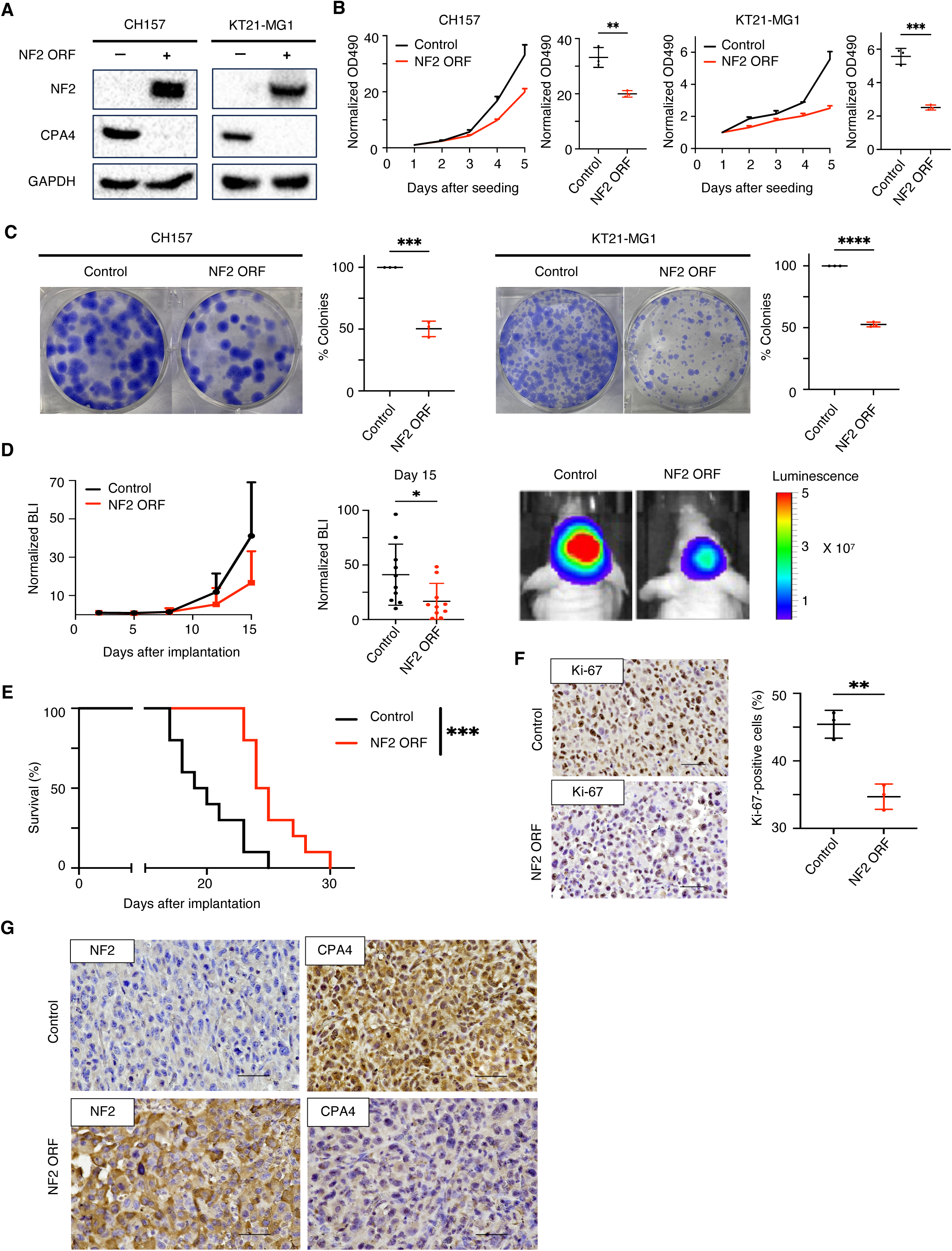
NF2 restoration suppresses CPA4 expression and NF2-deficient meningioma growth. (A) Immunoblot analysis of NF2 and CPA4 after control or NF2 ORF expression in CH157 and KT21- MG1 cells. (B) Growth curves and day-5 OD490 values. Data are mean ± SEM from three replicates normalized to day 1; unpaired, two-tailed t test: CH157, *P* = 0.0038; KT21-MG1, *P* = 0.0004. (C) Representative colony-formation assays and quantification. Data are mean ± SEM from three replicates; unpaired, two-tailed t test: CH157, *P* = 0.0002; KT21-MG1, *P* < 0.0001. (D) Growth curves, day-15 BLI values, and representative images of orthotopic CH157 xenografts expressing control or NF2 ORF (n = 10 per group). BLI values are mean ± SD; unpaired, two-tailed t test, *P* = 0.0286. (E) Kaplan–Meier survival analysis; log-rank test, *P* = 0.0009. (F) Ki-67 staining and quantification in intracranial tumors. Data are mean ± SEM from three tumors; unpaired, two-tailed t test, *P* = 0.0026. (G) NF2 and CPA4 IHC in intracranial tumors. Magnification, 400× (40× objective).

### CPA4 expression is enriched in NF2-deficient meningioma and associates with aggressive clinicogenomic features

To define the relationship between *NF2* and *CPA4* expression in human meningiomas, we interrogated meningioma datasets in the R2: Genomics Analysis and Visualization Platform (19). Across the dataset, *NF2* and *CPA4* expression showed a strong inverse correlation (Figure 3A). *CPA4* expression was significantly higher in *NF2*-mutant than in *NF2*-wild-type meningiomas and was likewise elevated in tumors with chromosome 22q loss, a recurrent alteration that encompasses NF2 (Figure 3B) (20). We next examined the relationship between *CPA4* expression and clinicogenomic features using integrated meningioma datasets in Oncoscape (6, 7, 21–28). WHO grade and chromosome 1p status, an established marker of aggressive meningioma biology (2), were available for analysis. Tumors were stratified into CPA4-low and CPA4-high groups. Grade 3 and grade 2 tumors comprised 5.2% and 25.8% of the CPA4-low group, respectively, compared with 7.6% and 35.6% of the CPA4-high group (Figure 3C). Chromosome 1p loss was also more frequent in CPA4-high than in CPA4-low tumors (41.4% versus 20.3%; Figure 3C). Thus, elevated *CPA4* expression associates with higher WHO grade and chromosome 1p loss in meningioma.

**Figure 3.**
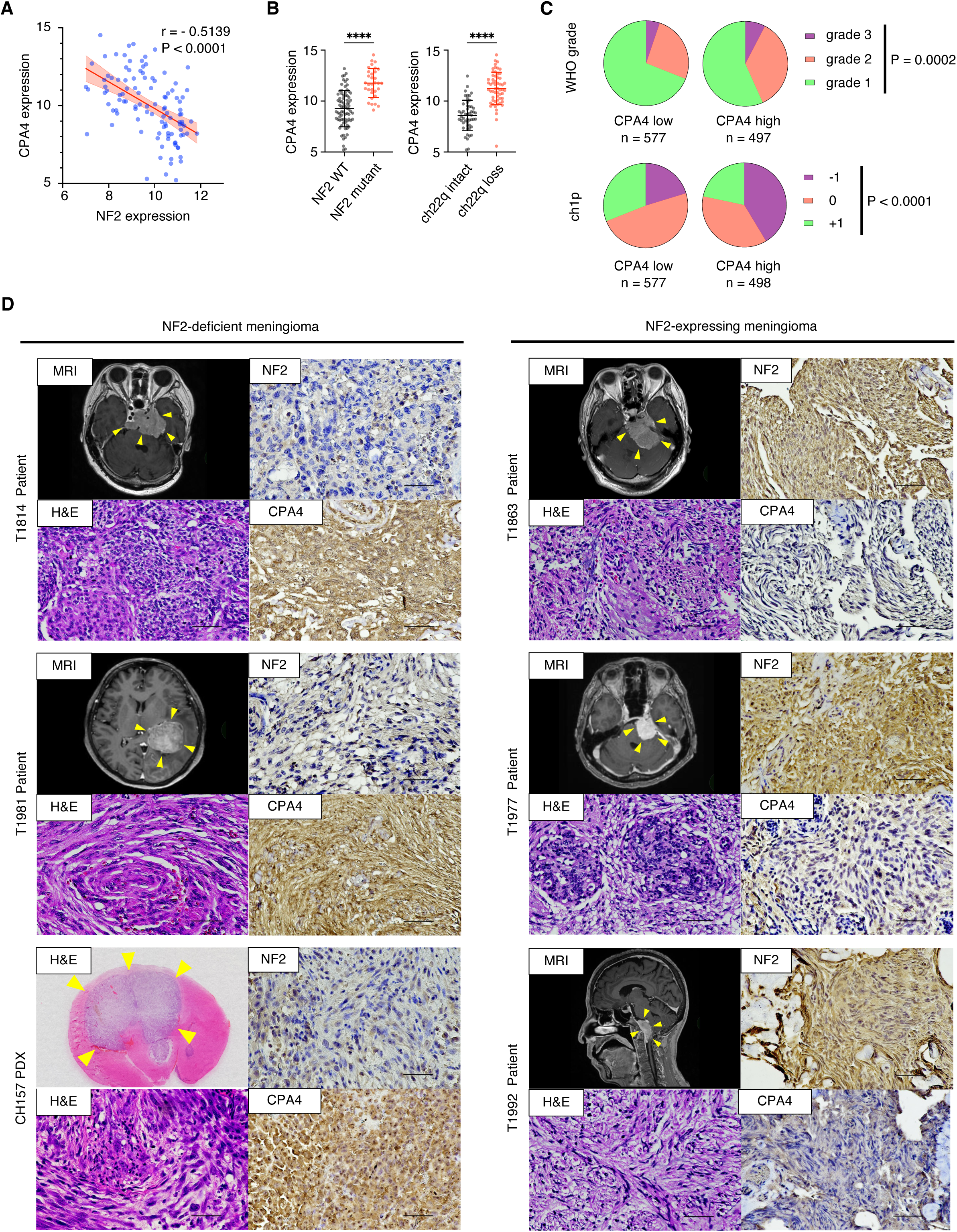
CPA4 is enriched in NF2-deficient meningioma and associates with aggressive clinicogenomic features. (A) Correlation between NF2 and CPA4 expression in meningiomas from R2 (n = 109); Pearson r = −0.5139, *P* < 0.0001. (B) CPA4 expression in NF2-wild-type (n = 77) versus NF2- mutant (n = 32) meningiomas and in tumors with intact (n = 51) versus deleted (n = 58) chromosome 22q. Unpaired, two-tailed t tests, *P* < 0.0001 for both comparisons. (C) WHO grade and chromosome 1p status in Oncoscape meningiomas stratified by CPA4 expression (CPA4-low, n = 578; CPA4-high, n = 500). Cases lacking the relevant annotation were excluded. Chi-square tests: WHO grade, *P* = 0.0002; chromosome 1p status, *P* < 0.0001. (D) MRI, H&E, and NF2 and CPA4 IHC in orthotopic CH157 xenografts and patient meningiomas. Magnification, 400× (40× objective).

We next evaluated NF2 and CPA4 protein expression by IHC in five patient meningiomas (T1814, T1981, T1863, T1977, and T1992) and orthotopic CH157 xenografts. Consistent with the public-dataset analyses, NF2-deficient patient tumors (T1814 and T1981) and CH157 xenografts exhibited high *CPA4* expression, whereas NF2-expressing patient tumors (T1863, T1977, and T1992) showed low *CPA4* expression (Figure 3D).

Given the frequent loss-of-function alterations in NF2 in mesothelioma (10, 29, 30), we analyzed *CPA4* transcript levels across The Cancer Genome Atlas (TCGA) pan-cancer datasets through UCSC Xena (31) to determine whether *CPA4* elevation extends beyond meningioma. *CPA4* expression was among the highest in mesothelioma (Supplementary Figure S1A). Mesotheliomas with high *CPA4* expression had a shorter median overall survival than those with low *CPA4* expression (457 versus 630 days), although the difference was not statistically significant (*P* = 0.1384; Supplementary Figure S1B). These exploratory observations identify mesothelioma as an additional NF2-altered tumor context characterized by high *CPA4* expression.

### CPA4 is required for NF2-deficient meningioma growth and tumor maintenance

To directly test whether CPA4 represents a functional dependency in NF2-deficient meningioma, we depleted CPA4 in CH157 and KT21-MG1 cells using a doxycycline (dox)-inducible shRNA system that enabled controlled and reversible target suppression. The vector design is shown in Supplementary Figure 2A, and induction of the shRNA construct was verified by red fluorescence, which was extinguished following dox withdrawal (Supplementary Figure S2B). Immunoblotting confirmed efficient CPA4 protein depletion during dox exposure and restoration of CPA4 expression after dox removal (Figure 4A). CPA4 depletion significantly reduced cell proliferation and colony formation in both NF2-deficient meningioma models (Figure 4B and C). Doxycycline withdrawal restored proliferation and clonogenic growth toward untreated control levels, linking the growth phenotype to sustained CPA4 depletion. These findings identify CPA4 as a reversible dependency required to maintain NF2-deficient meningioma cell growth in vitro.

**Figure 4.**
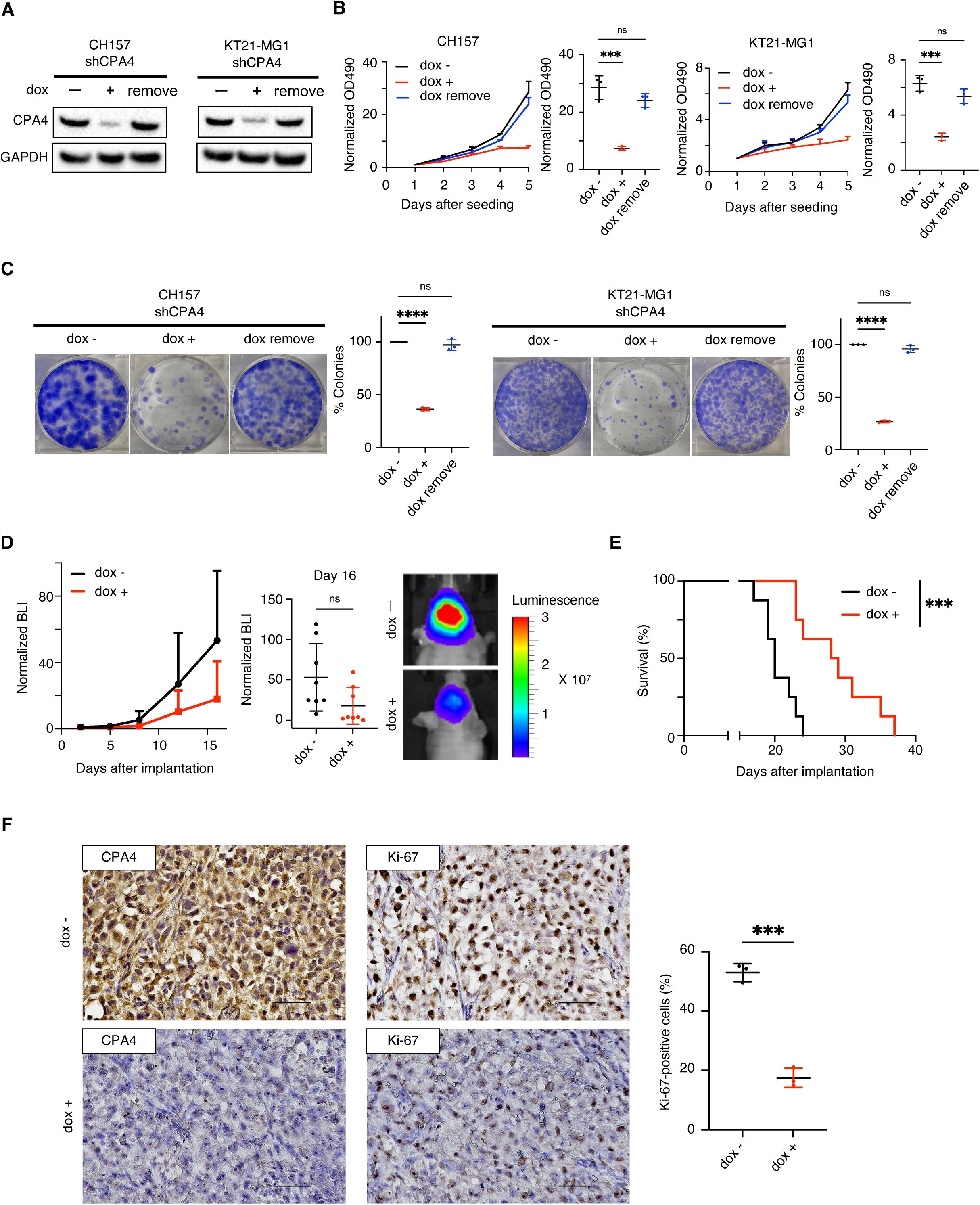
CPA4 is required to sustain NF2-deficient meningioma growth and tumor maintenance. (A) Immunoblot analysis showing doxycycline-inducible CPA4 depletion and restoration after doxycycline withdrawal in CH157 and KT21-MG1 cells. (B) Growth curves and day-5 OD490 values showing reversible growth suppression. Data are mean ± SEM from three replicates normalized to day 1. One-way ANOVA with Tukey test: CH157, doxycycline− versus doxycycline+, *P* = 0.0002; doxycycline− versus doxycycline withdrawal, *P* = 0.1982; KT21-MG1, doxycycline− versus doxycycline+, *P* = 0.0001; doxycycline− versus doxycycline withdrawal, *P* = 0.1131. (C) Colony-formation assays and quantification. Data are mean ± SEM from three replicates. One-way ANOVA with Tukey test: CH157, doxycycline− versus doxycycline+, *P* < 0.0001; doxycycline− versus doxycycline withdrawal, *P* = 0.5563; KT21-MG1, doxycycline− versus doxycycline+, *P* < 0.0001; doxycycline− versus doxycycline withdrawal, *P* = 0.1081. (D) Growth curves, day-16 BLI values, and representative images from orthotopic CH157 xenografts expressing doxycycline-inducible CPA4 shRNA (vehicle, n = 8; doxycycline, n = 8). Data are mean ± SD; unpaired, two-tailed t test, *P* = 0.0553. (E) Kaplan–Meier survival analysis; log-rank test, *P* = 0.0008. (F) CPA4 and Ki-67 IHC and Ki-67 quantification. Data are mean ± SEM from three tumors; unpaired, two-tailed t test, *P* = 0.0002. Magnification, 400× (40× objective).

We next evaluated whether CPA4 was also required for tumor maintenance in vivo. CH157 cells carrying dox-inducible CPA4 shRNA were orthotopically implanted in mice, which then received daily intraperitoneal dox or vehicle. CPA4 depletion attenuated the trajectory of intracranial tumor growth measured by BLI, although the difference in tumor burden at day 16 did not reach statistical significance (*P* = 0.0553), and significantly prolonged survival (*P* = 0.0008; Figure 4D and E). IHC analysis confirmed reduced CPA4 expression in dox-treated tumors and demonstrated a marked decrease in Ki-67-positive proliferating cells (*P* = 0.0002; Figure 4F). Together, the reversible in vitro phenotype and the survival and pharmacodynamic effects observed in vivo establish CPA4 as a functional dependency that sustains NF2-deficient meningioma growth.

### CPA4 depletion reprograms proliferative and genome-maintenance pathways in NF2- deficient meningioma

To define the transcriptional programs maintained by CPA4 in NF2-deficient meningioma cells, we performed RNA sequencing in CH157 cells following inducible *CPA4* depletion. RNA-sequencing analysis confirmed robust suppression of *CPA4* transcripts (Figure 5A). Principal component analysis of the 5,000 most variable genes separated CPA4-depleted cells from controls along the first principal component, which accounted for 91% of the variance, demonstrating a broad CPA4-dependent transcriptional state (Supplementary Figure S3A). Differential-expression analysis identified 3,886 significantly downregulated and 3,551 significantly upregulated genes following CPA4 depletion (Figure 5B). Heatmap analysis further demonstrated coordinated suppression of genes governing cell-cycle progression (*MYBL2*, *CDK1*, *TOP2A*, and *CCNA2*), DNA replication (*CDC45*, *MCM2*, *GINS1*, and *PCNA*), and DNA repair (*RAD51*, *ERCC8*, *DDB2*, and *RPA2*) (Figure 5C).

**Figure 5.**
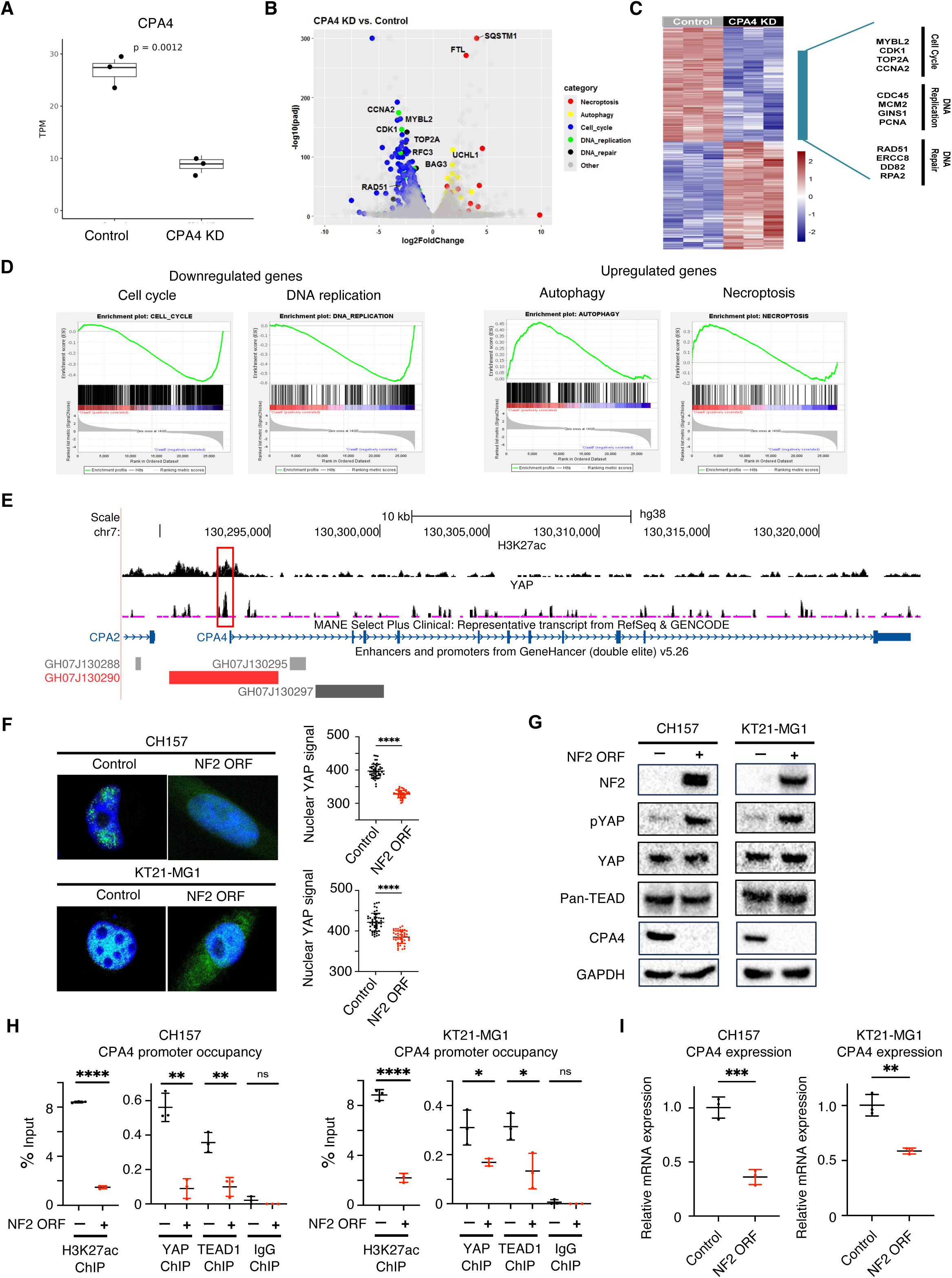
CPA4 sustains genome-maintenance programs and is directly regulated by YAP–TEAD in NF2-deficient meningioma. (A) CPA4 expression after inducible CPA4 depletion in CH157 cells. (B) Volcano plot of differentially expressed genes after CPA4 depletion, colored by biological program. (C) Heatmap highlighting coordinated changes in cell-cycle, DNA-replication, and DNA-repair genes. (D) GSEA showing negative enrichment of cell-cycle and DNA-replication programs and positive enrichment of autophagy and necroptosis programs (adjusted *P* < 0.001). (E) H3K27ac and YAP CUT&RUN tracks at the CPA4 locus in CH157 cells; the CPA4 promoter is boxed. (F) Representative YAP immunofluorescence and nuclear signal quantification after NF2 restoration. Data are mean ± SEM from 50 cells; unpaired, two-tailed t tests, *P* < 0.0001 in both models. (G) Immunoblot analysis of phosphorylated YAP, total YAP, pan-TEAD, CPA4, and GAPDH after NF2 restoration. (H) ChIP-qPCR for H3K27ac, YAP, and TEAD1 at the CPA4 promoter after NF2 restoration. Data are mean ± SEM from three replicates; unpaired, two-tailed t tests: CH157, H3K27ac *P* < 0.0001, YAP *P* = 0.0012, TEAD1 *P* = 0.0051; KT21-MG1, H3K27ac *P* < 0.0001, YAP *P* = 0.0279, TEAD1 *P* = 0.0258. (I) CPA4 mRNA after NF2 restoration. Data are mean ± SEM from three replicates; unpaired, two-tailed t tests: CH157, *P* = 0.0008; KT21-MG1, *P* = 0.0021.

Pathway analysis reinforced this pattern: cell-cycle, DNA-replication, homologous-recombination, and DNA-repair pathways were among the most strongly suppressed programs after CPA4 depletion (Supplementary Figure S3B). Gene set enrichment analysis further confirmed negative enrichment of cell-cycle and DNA-replication programs (Figure 5D). Conversely, CPA4 depletion induced genes linked to autophagy or necroptosis, including *BAG3*, *UCHL1*, *SQSTM1*, and *FTL* (Figure 5B and C), accompanied by positive enrichment of autophagy and necroptosis programs (Figure 5D; Supplementary Figure S3B). Thus, CPA4 depletion shifts the transcriptional state of NF2-deficient meningioma cells from proliferative and genome-maintenance programs toward stress and cell-death programs, defining a CPA4-dependent transcriptional program that supports tumor maintenance.

### CPA4 is a direct YAP–TEAD transcriptional target in NF2-deficient meningioma

Because NF2 deficiency activates nuclear YAP–TEAD signaling and is consistently associated with elevated CPA4 expression, we investigated whether CPA4 is directly regulated by this pathway. HOMER motif analysis of publicly available H3K27ac CUT&Tag data from CH157 cells (SRR29264002) identified significant enrichment of TEAD-binding motifs within H3K27ac-marked regulatory regions (Supplementary Table S2), implicating YAP–TEAD as a candidate regulator of CPA4. We therefore mapped H3K27ac and YAP occupancy by CUT&RUN in NF2-deficient CH157 cells. Both H3K27ac and YAP were enriched at the *CPA4* promoter, revealing direct YAP engagement at an active CPA4 regulatory locus (Figure 5E).

We next determined whether NF2 restoration attenuated this regulatory state. NF2 restoration markedly reduced nuclear YAP in both CH157 and KT21-MG1 cells (Figure 5F). Immunoblotting further showed increased inhibitory YAP phosphorylation without appreciable changes in total YAP or pan-TEAD abundance, accompanied by a marked reduction in CPA4 protein (Figure 5G). To define the corresponding chromatin changes at the *CPA4* promoter, we performed ChIP-qPCR for H3K27ac, YAP, and TEAD1 using primers spanning the H3K27ac- and YAP-enriched region identified by CUT&RUN. NF2 restoration significantly reduced H3K27ac enrichment and YAP and TEAD1 occupancy at the *CPA4* promoter in both meningioma models, whereas IgG controls remained at background levels (Figure 5H). Concordantly, NF2 restoration significantly reduced *CPA4* mRNA expression in both cell lines (Figure 5I). Together, these orthogonal epigenomic, biochemical, and transcriptional data establish CPA4 as a direct YAP–TEAD transcriptional target activated by NF2 deficiency, providing a mechanistic basis for the CPA4 dependency observed in NF2-deficient meningioma.

### CPA4 marks YAP–TEAD inhibitor sensitivity in NF2-deficient meningioma

We examined whether CPA4 expression marks sensitivity to YAP–TEAD inhibition, an emerging therapeutic vulnerability in NF2-deficient tumors (9–12). Consistent with Figure 3, immunoblotting showed high CPA4 expression in the NF2-deficient meningioma cell lines CH157, KT21-MG1, T1814, and T1981, whereas CPA4 expression was low in the NF2-expressing cell lines T1863, T1977, and T1992 (Figure 6A). We compared verteporfin, a repurposed photosensitizer reported to disrupt YAP–TEAD transcriptional activity (17, 18), and VT3989, a clinical-stage pan-TEAD autopalmitoylation inhibitor (9). CPA4-high cells were consistently more sensitive to both agents than CPA4-low cells (Figure 6B). Verteporfin IC₅₀ values ranged from 2.7 to 3.9 μM in CPA4-high cells but exceeded 10 μM in CPA4-low cells; VT3989 IC₅₀ values ranged from 13.8 to 25.6 μM in CPA4-high cells but exceeded 50 μM in CPA4-low cells. Both inhibitors also produced time-dependent suppression of proliferation and reduced colony formation in CPA4-high meningioma cells (Supplementary Figure S4A, B).

**Figure 6.**
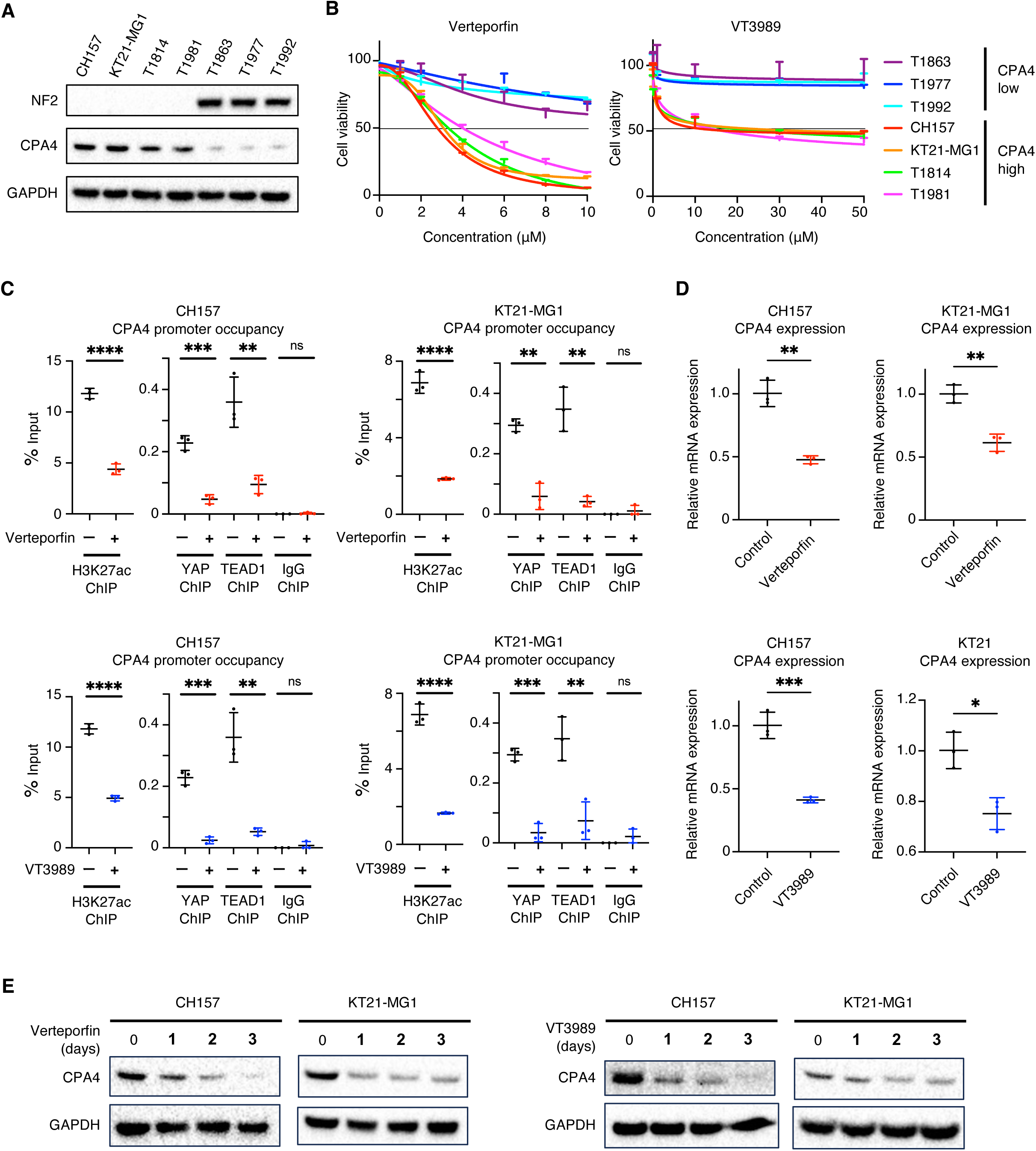
CPA4 expression marks sensitivity and pharmacodynamic response to YAP–TEAD inhibition. (A) Immunoblot analysis of NF2 and CPA4 in NF2-deficient/CPA4-high and NF2- expressing/CPA4-low meningioma cells. (B) Dose–response curves for verteporfin and VT3989. Data are mean ± SEM from three replicates. (C) ChIP-qPCR for H3K27ac, YAP, and TEAD1 at the CPA4 promoter after verteporfin or VT3989 treatment. Data are mean ± SEM from three replicates; unpaired, two-tailed t tests: CH157 verteporfin, H3K27ac *P* < 0.0001, YAP *P* = 0.0004, TEAD1 *P* = 0.0059; CH157 VT3989, H3K27ac *P* < 0.0001, YAP *P* = 0.0002, TEAD1 *P* = 0.0029; KT21-MG1 verteporfin, H3K27ac *P* < 0.0001, YAP *P* = 0.0011, TEAD1 *P* = 0.0021; KT21-MG1 VT3989, H3K27ac *P* < 0.0001, YAP *P* = 0.0003, TEAD1 *P* = 0.0079. (D) CPA4 mRNA after verteporfin or VT3989 treatment. Data are mean ± SEM from three replicates; unpaired, two-tailed t tests: CH157 verteporfin, *P* = 0.0011; CH157 VT3989, *P* = 0.0007; KT21-MG1 verteporfin, *P* = 0.0025; KT21-MG1 VT3989, *P* = 0.0107. (E) Immunoblot analysis of CPA4 during treatment with verteporfin or VT3989 at the corresponding IC_50_ concentration.

We next determined whether pharmacologic YAP–TEAD inhibition suppressed *CPA4* transcriptional regulation. In both models, verteporfin and VT3989 significantly reduced H3K27ac enrichment and YAP and TEAD1 occupancy at the *CPA4* promoter, whereas IgG controls remained at background levels (Figure 6C). Both agents significantly reduced *CPA4* mRNA expression (Figure 6D) and produced a time-dependent decrease in CPA4 protein abundance (Figure 6E). Together, these data support CPA4 expression as a candidate response marker and pharmacodynamic readout of YAP–TEAD pathway inhibition in NF2-deficient meningioma.

### Targeting the YAP–TEAD–CPA4 axis suppresses orthotopic NF2-deficient meningioma and prolongs survival

Given the growth-inhibitory effects of YAP–TEAD inhibition in vitro, we next evaluated its therapeutic activity in vivo. Dose-tolerability studies showed that verteporfin at 75 mg/kg and VT3989 at 100 mg/kg were well tolerated, with no treatment-associated decline in body weight; these doses were therefore selected for efficacy studies (Supplementary Figure S5A, B). We treated mice bearing orthotopic NF2-deficient CH157 meningioma xenografts with these regimens. Both agents significantly reduced intracranial tumor burden at day 15, as measured by BLI, and prolonged survival (Figure 7A, B). Tumors collected after treatment showed reduced CPA4 staining and a significant decrease in Ki-67-positive tumor cells, linking pathway inhibition to suppression of CPA4 expression and tumor-cell proliferation in vivo (Figure 7C). Together with the mechanistic findings above, these data support a model in which NF2 deficiency engages a therapeutically actionable YAP–TEAD–CPA4 axis that sustains meningioma growth (Figure 7D).

**Figure 7.**
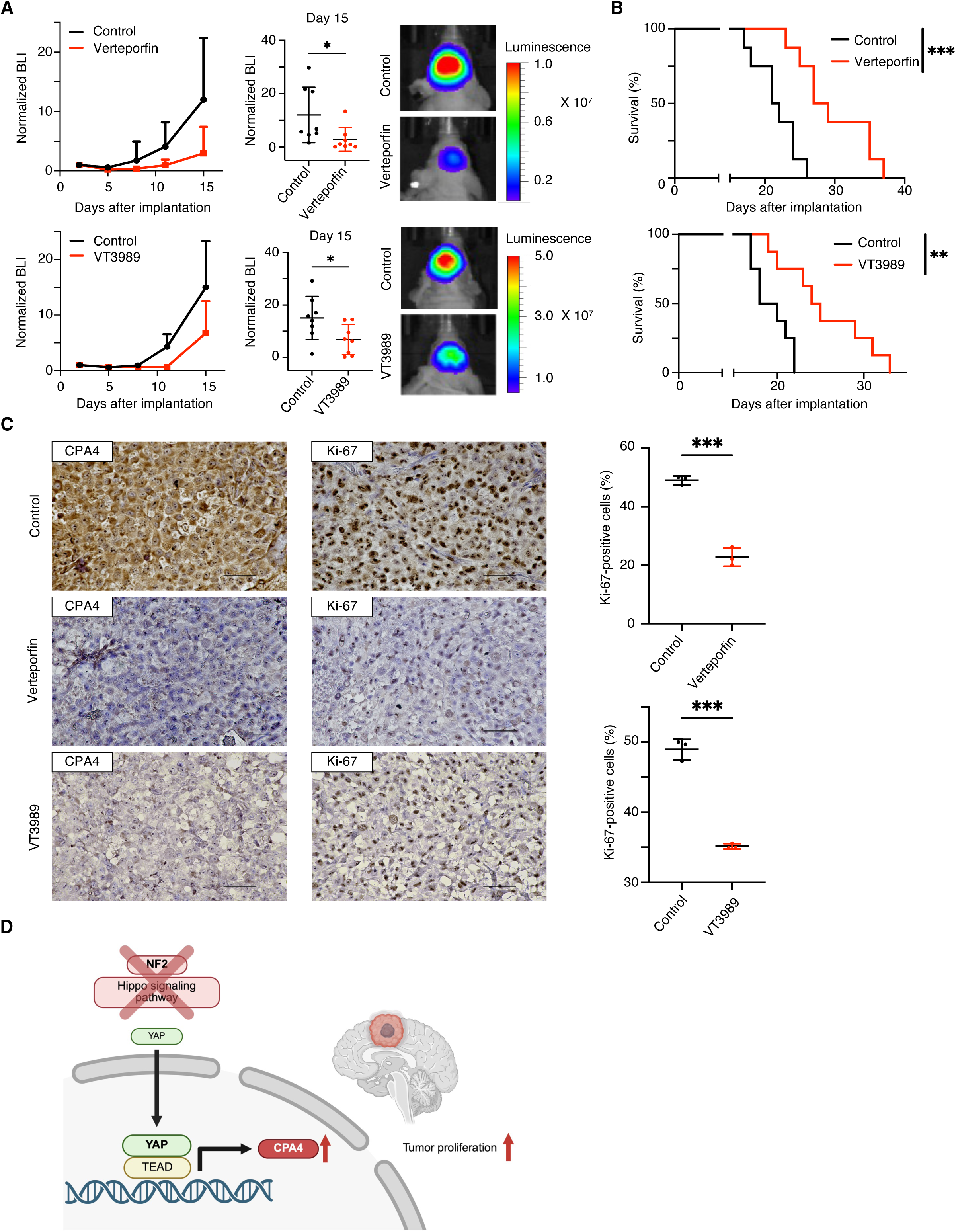
Targeting the YAP–TEAD–CPA4 axis suppresses orthotopic NF2-deficient meningioma and prolongs survival. Mice bearing orthotopic CH157 xenografts received vehicle or verteporfin (75 mg/kg intraperitoneally; n = 8 per group) or vehicle or VT3989 (100 mg/kg orally; n = 8 per group). (A) Tumor-growth curves, day-15 BLI values, and representative images. BLI values are normalized to treatment initiation and shown as mean ± SD; day-15 comparisons used unpaired, two-tailed t tests: verteporfin, *P* = 0.0393; VT3989, *P* = 0.0363. (B) Kaplan–Meier survival analyses; log-rank tests with Holm adjustment: verteporfin, *P* = 0.0008; VT3989, *P* = 0.0034. (C) CPA4 and Ki-67 IHC and Ki-67 quantification after treatment. Data are mean ± SEM from three tumors; unpaired, two-tailed t tests: verteporfin, *P* = 0.0002; VT3989, *P* = 0.0001. (D) Model of the NF2–YAP–TEAD–CPA4 axis in NF2- deficient meningioma.

## DISCUSSION

NF2 deficiency defines the most prevalent molecular driver of meningioma, yet its therapeutic implications remain incompletely resolved because the downstream effectors that convert YAP– TEAD activation into sustained tumor growth are poorly defined (3–12). Here, we uncover CPA4 as a direct YAP–TEAD transcriptional effector required to sustain NF2-deficient meningioma. CPA4 expression was consistently enriched in *NF2*-mutant and chromosome 22q-deleted meningiomas and associated with higher WHO grade and chromosome 1p loss. Genetic CPA4 depletion impaired tumor-cell proliferation, suppressed intracranial tumor growth, and prolonged survival, whereas CPA4-high meningioma cells showed preferential sensitivity to YAP–TEAD inhibition. Moreover, verteporfin and the clinical-stage TEAD inhibitor VT3989 reduced CPA4 expression, suppressed orthotopic tumor growth, and prolonged survival. These findings position CPA4 at the intersection of NF2 deficiency, oncogenic YAP–TEAD transcription, tumor maintenance, and therapeutic response and establish a targetable YAP–TEAD–CPA4 axis in NF2-deficient meningioma (Figure 7D).

Although CPA4 has been associated with tumor progression, treatment resistance, and aggressive clinical behavior in several cancer types (13–16), its mechanistic contribution to meningioma and functional relationship with NF2 signaling had not been defined. Our study establishes CPA4 as an active tumor-maintaining effector rather than a passive correlate of an adverse meningioma phenotype. CPA4 depletion reversibly suppressed proliferation and colony formation (Figure 4) and produced coordinated transcriptional attenuation of cell-cycle, DNA-replication, and DNA-repair programs, together with induction of autophagy and necroptosis programs (Figure 5A–D). The convergence of these molecular changes with the in vitro and in vivo growth phenotypes demonstrates that CPA4 sustains a broad tumor-maintenance state rather than exerting a limited effect on proliferation. Thus, the genetic and transcriptional evidence establishes CPA4 as a functional dependency of NF2-deficient meningioma and a potential therapeutic target.

The upstream mechanism controlling *CPA4* expression in cancer has also remained poorly defined. HIF-1α-dependent induction of *CPA4* expression has been reported under hypoxic conditions (32), but a direct connection between NF2 loss, YAP–TEAD activity, and *CPA4* transcription had not been established. Our study resolves this gap through multiple, orthogonal lines of epigenomic and functional evidence. TEAD motifs were enriched within active regulatory regions; YAP and H3K27ac occupied the *CPA4* promoter; and NF2 restoration reduced nuclear YAP, decreased YAP and TEAD1 occupancy at the *CPA4* promoter, and diminished H3K27ac enrichment and *CPA4* transcription. Pharmacologic YAP–TEAD inhibition reproduced these promoter-level and transcriptional effects (Figure 5E–I). These data identify CPA4 as a direct, dynamically regulated YAP–TEAD target rather than a secondary consequence of NF2-deficient tumor growth. The mechanism uncovered here also provides a unifying explanation for CPA4 elevation in *YAP1*–*MAML2*-positive meningiomas, which share YAP-related transcriptional profiles with *NF2*-mutant tumors (33), and for CPA4 suppression together with *CCN1*, *CCN2*, and *ANKRD1* following combined brigatinib and INK128 treatment in an NF2-associated meningioma model (34). The inclusion of *CPA4* in independently derived YAP–TEAD gene signatures further supports this relationship (35, 36). Thus, our study mechanistically connects previously separate observations and establishes *CPA4* as a recurrent transcriptional effector of oncogenic YAP– TEAD activity.

The translational significance of this mechanism is heightened by the clinical development of TEAD inhibitors. YAP–TEAD signaling is an established therapeutic vulnerability in NF2-deficient mesothelioma, schwannoma, and meningioma (10–12), and VT3989 has shown early clinical activity in advanced solid tumors enriched for NF2 alterations (9). Our findings address a central translational need by linking pathway activity and drug sensitivity to a measurable downstream factor. High baseline CPA4 expression marked preferential sensitivity to two mechanistically distinct inhibitors (Figure 6), while treatment reduced YAP and TEAD1 occupancy at the *CPA4* promoter and decreased *CPA4* mRNA and protein in vitro and in vivo (Figures 6C–E and 7C). The concordant effects of verteporfin and the selective TEAD autopalmitoylation inhibitor VT3989 strengthen the link between pathway inhibition and CPA4 suppression. CPA4 therefore has two complementary biomarker properties: baseline expression marks YAP–TEAD inhibitor sensitivity, and its on-treatment reduction reports pathway suppression. CPA4 may also complement genomic *NF2* status by capturing YAP–TEAD activation in *NF2*-wild-type tumors, including *YAP1*–*MAML2* fusion-positive meningiomas. Because CPA4 is secreted and circulating CPA4 has been detected in other cancers (13, 15), these properties provide a practical foundation for incorporating tumor and circulating CPA4 measurements into studies of TEAD-directed therapy. Pan-cancer analysis identified mesothelioma as an additional NF2-altered tumor type with high CPA4 expression, broadening the relevance of the YAP–TEAD–CPA4 relationship beyond meningioma (Supplementary Figure S1). Although the experimental models used here were derived from sporadic meningiomas, their shared NF2-deficient state supports the potential relevance of this pathway to NF2-associated tumors. The specific CPA4 substrates that connect its enzymatic activity to the transcriptional programs observed here remain unresolved; nevertheless, the convergence of patient-cohort analyses, genetic perturbation, orthogonal epigenomic assays, pharmacologic inhibition, and orthotopic efficacy establishes a coherent mechanistic and translational framework. Collectively, our study uncovers CPA4 as a tumor-maintaining effector of oncogenic YAP–TEAD signaling and establishes the YAP–TEAD–CPA4 axis as a biomarker-linked therapeutic vulnerability in NF2-deficient meningioma, with broader implications for NF2-altered cancers.

## MATERIALS AND METHODS

### Sex as a biological variable

Our study examined 6-week-old female athymic mice (rnu/rnu genotype, BALB/c background). The animals were purchased from Inotiv and housed under aseptic conditions. Meningiomas exhibit a female predominance in the general population (37), although the biological mechanisms underlying this sex disparity remain incompletely understood. NF2 is a well-known risk factor for meningioma development (37), but studies of individuals with NF2 have not demonstrated a consistent overall sex bias for meningioma incidence (38). In addition, molecularly defined NF2/22q-altered meningiomas may show a reduced female predominance compared with meningiomas overall (39, 40). The study was designed to investigate tumor-intrinsic NF2-deficient meningioma biology in a controlled in vivo context, and available data from NF2 and NF2-altered meningioma patients do not support a consistent sex-dependent influence on tumor incidence or behavior.

### Cell sources and propagation

CH157-MN cells (hereafter CH157; RRID: CVCL_5723) were obtained from an existing laboratory stock at the University of Alabama at Birmingham. KT21-MG1-Luc5 cells, derived from KT21- MG1 (RRID: CVCL_M429), were provided by Long-Sheng Chang (The Ohio State University, Columbus, OH, USA). KT21-MG1-Luc5 was derived from a spontaneously arising malignant meningioma and engineered to express luciferase and neomycin resistance. CH157 and KT21- MG1 were derived from female patients. T1814, T1981, T1863, T1977, and T1992 cells were provided by Yukitomo Ishi and Shigeru Yamaguchi (Hokkaido University, Sapporo, Japan). These five primary cultures were established from patient meningiomas, as shown by MRI in Figure 3D. CH157, T1814, T1981, T1863, T1977, and T1992 cells were propagated as monolayers in complete medium consisting of DMEM (Corning, 10013CV) supplemented with 10% FBS (Corning, 35010CV). KT21-MG1-Luc5 cells were propagated as monolayers in tumor stem medium (TSM) supplemented with 5% FBS. TSM base consisted of Neurobasal-A medium (Gibco, 10888022), DMEM/F-12 with GlutaMAX (Gibco, 10565018), HEPES (Gibco, 15630080), sodium pyruvate (Gibco, 11360070), MEM nonessential amino acids (Gibco, 11140050), B-27 supplement minus vitamin A (Gibco, 12587010), EGF and FGF (FUJIFILM Irvine Scientific, 10026 and 100146), PDGF-AA and PDGF-BB (FUJIFILM Irvine Scientific, 10016 and 10018), and 0.2% heparin (STEMCELL Technologies, 07980). Cell identity was confirmed by short tandem repeat profiling using the PowerPlex 16 HS System (Promega, DC2101). Cells were maintained at 37°C in a humidified atmosphere of 95% air and 5% CO₂ and tested negative for mycoplasma using the Mycoplasma Detection Kit (Abcam, ab289834).

### cDNA open reading frame (ORF) and short hairpin RNA (shRNA) treatments

Control ORF (Horizon Discovery, OHS5832), NF2 ORF (Horizon Discovery, PLOHS_100000483), and CPA4 shRNA (Horizon Discovery, V3THS_351959) were used to generate lentiviruses and infect tumor cells according to the manufacturer’s instructions.

### Cell viability assay

Tumor cells were seeded in 96-well plates at 2,000 cells per well and cultured in the presence of verteporfin (MedChemExpress, HY-B0146) or VT3989 (Selleckchem, E6486) for 72 hours with triplicate samples for each incubation condition. Relative numbers of viable cells were determined using CellTiter 96 AQueous One Solution Cell Proliferation Assay (Promega, G3581). IC₅₀ values were calculated using nonlinear least-squares curve-fitting. The proliferation effects by these inhibitors were analyzed on days 1, 2, 3, 4, and 5 in the presence of vehicle (0.5% DMSO) and IC₅₀ concentrations of verteporfin or VT3989. The proliferation effects by *NF2* overexpression and CPA4 dox-inducible shRNA were analyzed on days 1, 2, 3, 4, and 5. The effects by *NF2* overexpression were evaluated using cells transduced with control ORF as the control group and cells transduced with *NF2* ORF as the NF2-overexpressed group. The effects of CPA4 dox-inducible shRNA were evaluated using cells without doxycycline as the control group, cells treated with doxycycline for 96 hours as the CPA4 shRNA-expressing group, and cells 96 hours after doxycycline withdrawal as the doxycycline removal group.

### Colony formation assay

Colony formation assays were performed by plating 2,000 cells per well in 6-well plates. The cells were allowed to adhere for 24 hours and were then treated with 0.5% DMSO or IC₅₀ concentrations of verteporfin or VT3989. Cells were incubated at 37°C for two weeks, and colonies were counted following staining with 0.05% crystal violet. Cells modified with control ORF, *NF2* ORF, or CPA4 shRNA were also seeded at 2,000 cells per well in 6-well plates and incubated for two weeks. The effects by *NF2* overexpression were evaluated using cells transduced with control ORF as the control group and cells transduced with *NF2* ORF as the NF2-overexpressed group. The effects of CPA4 dox-inducible shRNA on colony formation were evaluated using cells without doxycycline as the control group, cells treated with doxycycline for 96 hours as the CPA4 shRNA-expressing group, and cells 96 hours after doxycycline withdrawal as the doxycycline removal group.

### Immunoblotting

Cells were cultured and treated with control ORF, NF2 ORF, CPA4 shRNA, verteporfin, or VT3989 (IC₅₀ concentrations). Total cell lysate was collected from the cell pellets at each condition using lysis buffer (Thermo Scientific, 89900) supplemented with a 1% protease and phosphatase inhibitor cocktail (Thermo Scientific, 78440), resolved by sodium dodecyl sulfate polyacrylamide gel electrophoresis, and then immunoblotted using standard techniques. Transfer was conducted using 1× NuPAGE transfer buffer (Invitrogen, NP00061) with 10% methanol. The protein samples were transferred to PVDF Transfer Membranes (Thermo Scientific, 88520) for 90 minutes at 30 V. The membranes were blocked for 1 hour at room temperature in 5% skim milk in Tris-HCl buffered saline with 0.05% Tween 20 (TBST), and then probed with primary antibodies (1:1000 dilution) overnight at 4°C. The membranes were then washed three times in TBST, incubated with secondary antibody for 1 hour (1:1000 dilution), and washed in TBST. Signals were visualized by enhanced chemiluminescence (Thermo Scientific, 32106). The primary antibodies used were NF2 (Cell Signaling Technology, 6995, RRID: AB_10828709), GAPDH (Cell Signaling Technology, 3683, RRID: AB_1642205), CPA4 (Proteintech, 26824-1-AP, RRID: AB_2880648), Phospho-YAP (Ser127) (Cell Signaling Technology, 13008, RRID: AB_2650553), YAP (Cell Signaling Technology, 14074, RRID: AB_2650491), and Pan-TEAD (Cell Signaling Technology, 13295, RRID: AB_2687902). The secondary antibody used was Anti-Rabbit IgG (Cell Signaling Technology, 7074, RRID: AB_2099233).

### Immunocytochemistry

Cells were seeded at 50,000 cells per well in 12-well plates containing coverslips. The cells were allowed to adhere for 24 hours. 4% paraformaldehyde (Santa Cruz Biotechnology, SC281692) was used to fix cells on coverslips. Slips were then rinsed three times in PBS and treated with 0.1% Triton X-100 for 15 minutes, followed by a blocking step in 5% bovine serum albumin (BSA) in PBS for 30 minutes at room temperature. Coverslips were incubated overnight at 4°C with YAP antibody (Cell Signaling Technology, 14074; RRID: AB_2650491) at a 1:200 dilution in PBS containing 5% BSA. Coverslips were washed in PBS three times and were incubated for 1 hour at room temperature in the dark with anti-rabbit antibody Alexa Fluor 488 (Cell Signaling Technology, 4412, RRID: AB_1904025) at 1:1000 dilution in PBS with 5% BSA for single reporter labeling. The coverslips were rinsed in PBS three times, and nuclei were stained and mounted on glass slides using Vectashield with DAPI (Vector Laboratories, H18002). Slides were analyzed with the Nikon Eclipse Ts2 microscope (Nikon), using the regions of interest (ROI) tool in the NIS-Elements software to measure GFP signals in cell nuclei. 50 nuclei per treatment condition were analyzed.

### RT-qPCR

RNA was extracted from cultured cell pellets using the RNeasy Mini Kit (Qiagen, 74104). RNA samples were treated with DNase I (Ambion, AM2222) prior to reverse transcription using RevertAid First Strand cDNA Synthesis Kit (Thermo Scientific, K1621). *HPRT1* and *CPA4* primers for RT-qPCR were obtained from PrimerBank (ID: 164518913c1 and 254540195c2, respectively). qPCR was performed using PowerUp SYBR Green Master Mix (Applied Biosystems, A25742) on a CFX96 Real-Time System with C1000 Touch Thermal Cycler (Bio-Rad). All reactions were performed in triplicate. The Ct value of target mRNA was normalized against that of *HPRT1* (ΔCt = Ct_target − Ct_*HPRT1*), and relative mRNA expression was calculated using the 2^−ΔΔCt method (ΔΔCt = ΔCt_treated − ΔCt_control). The primer sequences were as follows: *HPRT1*- forward: CCTGGCGTCGTGATTAGTGAT, *HPRT1*-reverse: AGACGTTCAGTCCTGTCCATAA, *CPA4*-forward: ATGGAGACGAGATCAGCAAATTG, *CPA4*-reverse: CAGGCCGATTGAAGGAGGAG. The effects by *NF2* overexpression were evaluated using cells transduced with control ORF as the control group and cells transduced with *NF2* ORF as the NF2- overexpressed group. The effects by YAP-TEAD inhibitors were analyzed in the presence of vehicle (0.5% DMSO) and IC₅₀ concentrations of verteporfin or VT3989 for 48 hours.

### ChIP-qPCR

Chromatin immunoprecipitation (ChIP) was performed using the SimpleChIP Enzymatic Chromatin IP Kit (Cell Signaling Technology, 9003) according to the manufacturer’s instructions. Immunoprecipitation was performed using the following antibodies: YAP (Cell Signaling Technology, 14074, RRID: AB_2650491), TEAD1 (Abcam, ab133533, RRID: AB_2737294), H3K27ac (Cell Signaling Technology, 8173, RRID: AB_10949503) and normal rabbit IgG (Cell Signaling Technology, 2729, RRID: AB_1031062) as a negative control. Two percent of the total chromatin was reserved as input. ChIP-qPCR enrichment was calculated as percent input (% Input) using the following formula: % Input = 2^(Ct_Input_adjusted − Ct_ChIP) × 100, where Ct_Input_adjusted = Ct_Input − log2(100/2), accounting for the 2% input fraction used. Results are presented as % Input for YAP, TEAD1, H3K27ac, and IgG ChIP at the target region. Primers for ChIP-qPCR were designed to target the YAP binding site within the CPA4 promoter, using NCBI Primer-BLAST. The target site was identified as follows: YAP ChIP-seq data from MDA-MB-231 cells (GSE66081) (41) were analyzed to extract YAP-enriched regions near the CPA4 locus. Genomic coordinates were converted from hg19 to hg38 using the UCSC LiftOver tool. Regions showing clustered YAP ChIP-seq reads were cross-referenced with our CUT&RUN (H3K27ac and YAP) data to identify accessible, transcriptionally active chromatin. The selected target region is located at chr7:130,292,767–130,293,131 (hg38), spanning the transcription start site of CPA4. The primer sequences were as follows: CPA4-forward: ATGCCAGGTTTATAATCTGACCTCT, CPA4-reverse: CTCCAGGAGCTGAAAAAGTCTG. The qPCR procedure and cell treatment conditions were the same as those used for RT-qPCR.

### RNA-sequencing analysis

RNA was extracted using the RNeasy Mini Kit (QIAGEN, 74104) according to the manufacturer’s instructions. Paired-end 150-bp FASTQ files were generated by Novogene and processed using the nf-core/rnaseq pipeline v3.1.0 (42) executed with Nextflow. Read quality and adapter content were assessed and trimmed within the pipeline using FastQC (RRID: SCR_014583) and Trim Galore!. Transcript-level abundance was quantified with Salmon v1.9.0 (43) against GRCh38. Gene-level counts were generated from quant.sf files using tximport, and differential expression was analyzed with DESeq2 (RRID: SCR_000154) (44) using default settings. Genes with a Benjamini–Hochberg-adjusted *P* value < 0.05 were considered differentially expressed. The DESeq2 Wald statistic was used to rank genes for gene set enrichment analysis with fgsea (Bioconductor) and MSigDB v2023.1 using 1,000 permutations, a maximum gene-set size of 500, and a minimum size of 20. DESeq2-normalized values were imported into GSEA v4.2.3 (RRID: SCR_003199) (45) to generate enrichment plots. Gene Ontology enrichment analysis was performed using ShinyGO v0.76.2 (46). Heatmaps were generated with the pheatmap R package; principal component, box, and volcano plots were generated in R using ggplot2 (RRID: SCR_014601).

### Cleavage under targets and release using nuclease (CUT&RUN)

Each reaction used 5 × 10^5 cells. CUT&RUN was performed with the CUTANA ChIC/CUT&RUN Kit (EpiCypher, 14-1048) according to the manufacturer’s protocol. Concanavalin A beads (11 μL per reaction) were activated, washed twice, resuspended, and combined with cells in wash buffer. After a 10-minute incubation, chilled antibody buffer was added. YAP (Cell Signaling Technology, 14074; RRID: AB_2650491) and H3K27ac (Cell Signaling Technology, 8173; RRID: AB_10949503) antibodies were used at 1:50, and samples were incubated overnight at 4°C. Samples were washed twice in cell-permeabilization buffer, incubated with pAG-MNase for 10 minutes at room temperature, washed twice, and resuspended in permeabilization buffer. Digestion was initiated with CaCl2 and continued for 2 hours at 4°C before addition of stop buffer and DNA purification. Libraries were prepared using the NEBNext Ultra II DNA Library Prep Kit and sequenced by MedGenome on a NovaSeq platform (paired-end, 100 bp). The nf-core/cutandrun pipeline (47) was used for adapter and quality trimming, alignment to GRCh38, counts-per-million normalization, and SEACR peak calling for CUT&RUN and public CUT&Tag data (SRR29264002). Peaks were annotated with ChIPseeker v1.22.1 (48).

### Xenograft studies

For orthotopic xenograft studies, mice received 3 μL of CH157 cell suspension (100,000 cells/μL) injected beneath the dura mater at the cerebral convexity, 3 mm lateral to bregma. For studies of NF2 restoration, mice were assigned to control ORF (n = 10) or *NF2* ORF (n = 10) groups. For CPA4-depletion studies, mice bearing CH157 tumors expressing doxycycline-inducible CPA4 shRNA received vehicle (n = 8) or doxycycline (n = 8) by daily intraperitoneal injection from treatment initiation until study endpoint. For verteporfin studies, mice received vehicle (n = 8) or verteporfin (75 mg/kg, intraperitoneally; n = 8). Mice were stratified by baseline bioluminescence signal and randomly assigned to treatment groups to ensure comparable tumor burden across groups. For VT3989 studies, mice received vehicle (n = 8) or VT3989 (100 mg/kg, orally; n = 8). Verteporfin and VT3989 were administered 5 days per week for 2 consecutive weeks. Tumor growth and treatment response were monitored by twice-weekly BLI. Mice were assessed daily and euthanized upon an irreversible neurologic deficit or a body condition score < 2. Mice reaching a prespecified humane endpoint were euthanized by CO₂ inhalation and cervical dislocation, in accordance with the approved IACUC protocol.

### Immunohistochemistry

FFPE slides from meningioma patients were provided by Yukitomo Ishi and Shigeru Yamaguchi (Hokkaido University, Sapporo, Japan). Brains were collected from the mice at 3 hours following completion of the last treatment (n = 3 for each treatment). Formalin-fixed brains were paraffin-embedded and sectioned for staining. FFPE slides from patient samples were stained using an H&E staining kit (Abcam, ab245880). The brain sections were stained with primary antibodies of NF2 (Proteintech, 21686-1-AP, RRID: AB_10792410), CPA4 (Proteintech, 26824-1-AP, RRID: AB_2880648), and anti-Ki-67 (Abcam, ab15580, RRID: AB_443209).

### Public data analysis

Publicly available data were accessed through the R2: Genomics Analysis and Visualization Platform (https://r2.amc.nl/), Oncoscape (https://oncoscape.sttrcancer.org/), and UCSC Xena (https://xenabrowser.net/) and analyzed using GraphPad Prism v10. Oncoscape analyses incorporated datasets from Fred Hutch Cancer Center (GEO: GSE252291) (6), University of California San Francisco (2018; GEO: GSE101638) (22), Baylor College of Medicine (GEO: GSE136661) (23), University of California San Diego (GEO: GSE139652) (24), University of California San Francisco (2020; GEO: GSE151921) (25), University of California San Francisco (2022; GEO: GSE183656) (7), Yale University School of Medicine (GEO: GSE85133) (26), Palacký University and University Hospital Olomouc (NIH BioProject: PRJNA705586) (27), and University of California San Francisco and University of Hong Kong (GEO: GSE212666) (28). Cases lacking WHO grade or chromosome 1p status were excluded only from the corresponding Oncoscape analysis. For TCGA analyses through UCSC Xena, cases without CPA4 expression data were excluded.

### Schematic model

The schematic model was created with BioRender (RRID: SCR_018361).

### Statistical analysis

Statistical analyses were performed using GraphPad Prism v10 (RRID: SCR_002798) unless otherwise specified. All statistical tests were two-sided, and *P* < 0.05 was considered statistically significant. Data are presented as mean ± SD or mean ± SEM, as indicated in the figure legends. The number of independent biological samples or animals (n) is specified in the corresponding legend. Two-group comparisons were performed using an unpaired two-tailed t test. Comparisons among three or more groups used one-way ANOVA followed by Tukey multiple-comparisons testing. Survival distributions were estimated by the Kaplan–Meier method and compared using the log-rank test with Holm adjustment when multiple survival comparisons were made. Associations of *CPA4* expression with WHO grade or chromosome 1p status were assessed using chi-square tests of independence. Correlations between *NF2* and *CPA4* expression were evaluated using Pearson correlation with a two-sided *P* value. Exact *P* values are reported when available; *P* < 0.0001 is reported for smaller values.

### Study approval

All animal protocols were approved by the University of Alabama at Birmingham Institutional Animal Care and Use Committee (protocol no. IACUC-22819). Human tissue specimens and associated MRI data were obtained under protocols approved by the Ethics Committee of Hokkaido University (approval no. 023-0199), with informed consent obtained from participants.

## Data availability

RNA-sequencing and CUT&RUN data generated in this study will be deposited in the NCBI Gene Expression Omnibus; accession numbers will be provided before publication. Public datasets and accession numbers used in this study are specified in Materials and Methods. Other data supporting the findings are available within the article, its Supplementary Data, and the Supporting Data Values file, or from the corresponding author upon reasonable request.

## AUTHOR CONTRIBUTIONS

KM, KP, and RH designed the study. KM and KP performed most of the experiments, and KM, KP, and RH wrote the manuscript. LC generated luciferase-expressing meningioma cells and edited the manuscript. KM, KP, NS, and RH performed and analyzed the in vitro experiments. KM, KP, RO, YM, YK, EU, DH, and RH performed and analyzed the in vivo experiments. KM, KP, and RH performed the public database analysis. SG performed all bioinformatics analyses and provided interpretation of the data. All authors commented on the manuscript and approved the included data.

## ACKNOWLEDGMENTS

We wish to acknowledge the Genomics Core Laboratory at the University of Alabama at Birmingham.

---

**Supplementary Figure S1.**
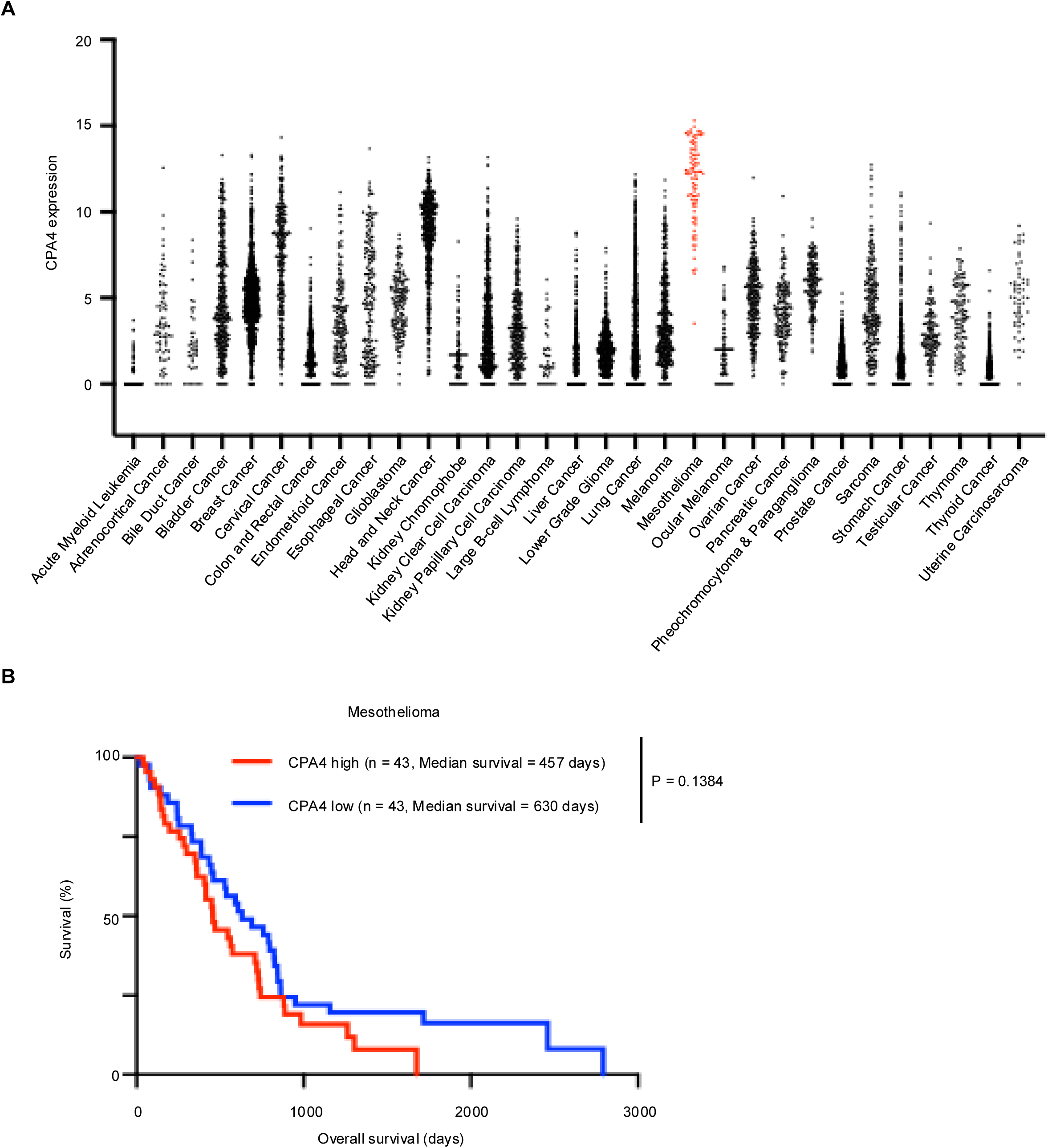
CPA4 expression in mesothelioma. (A) CPA4 expression across TCGA tumor types; mesothelioma is highlighted. (B) Overall survival of CPA4-high (n = 43) and CPA4-low (n = 43) TCGA mesotheliomas; log-rank test, *P* = 0.1384.

**Supplementary Figure S2.**
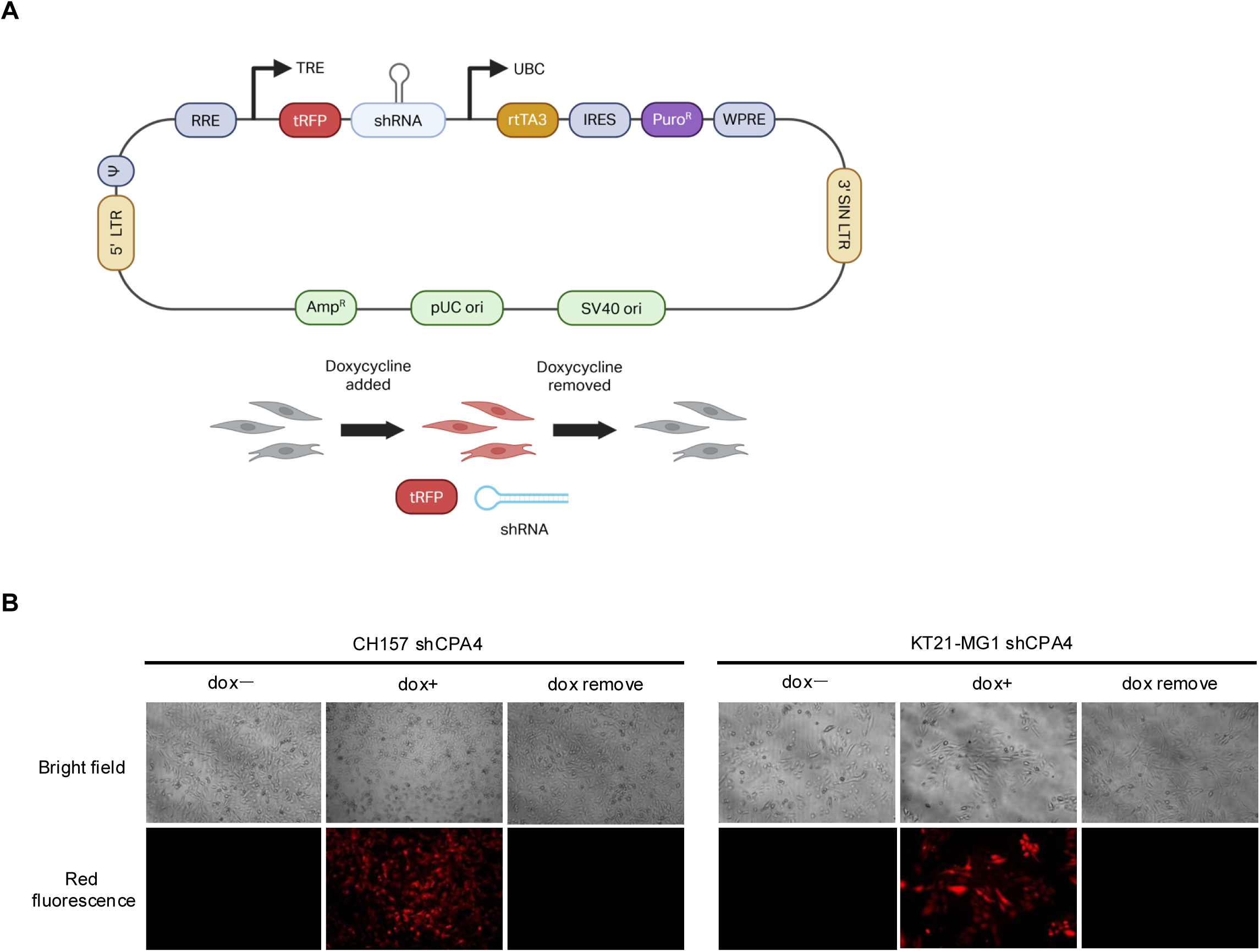
Doxycycline-inducible CPA4 depletion is reversible in NF2-deficient meningioma cells. (A) Doxycycline-inducible CPA4 shRNA vector and experimental design. (B) Bright- field and red-fluorescence images showing shRNA induction after doxycycline exposure and loss after withdrawal in CH157 and KT21-MG1 cells. Magnification, 100× (10× objective).

**Supplementary Figure S3.**
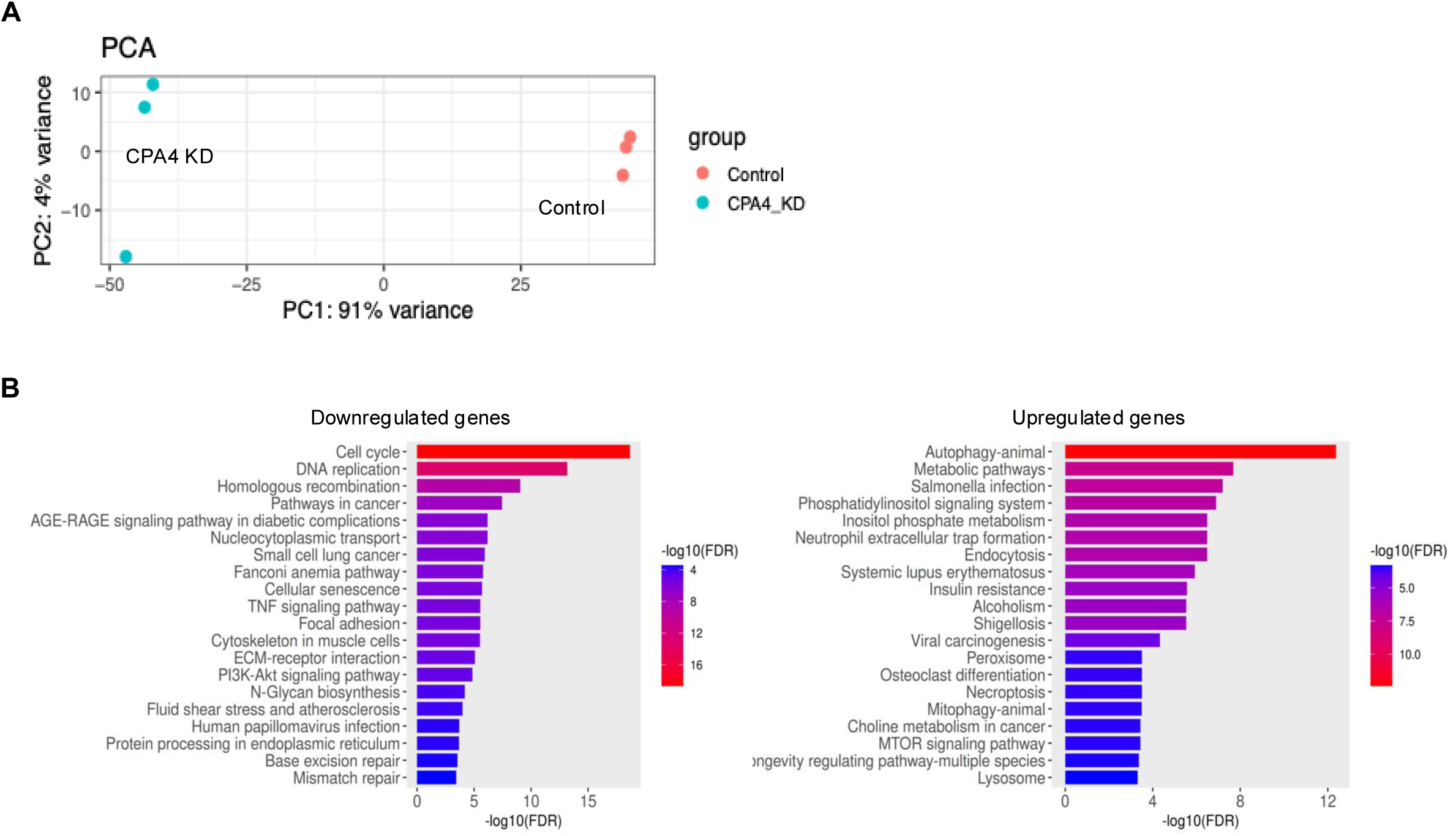
CPA4 depletion reprograms proliferative and genome-maintenance pathways. (A) Principal component analysis of RNA-sequencing profiles from control and CPA4- depleted CH157 cells (three biological replicates per group). (B) Gene Ontology enrichment analysis of downregulated and upregulated pathways after CPA4 depletion.

**Supplementary Figure S4.**
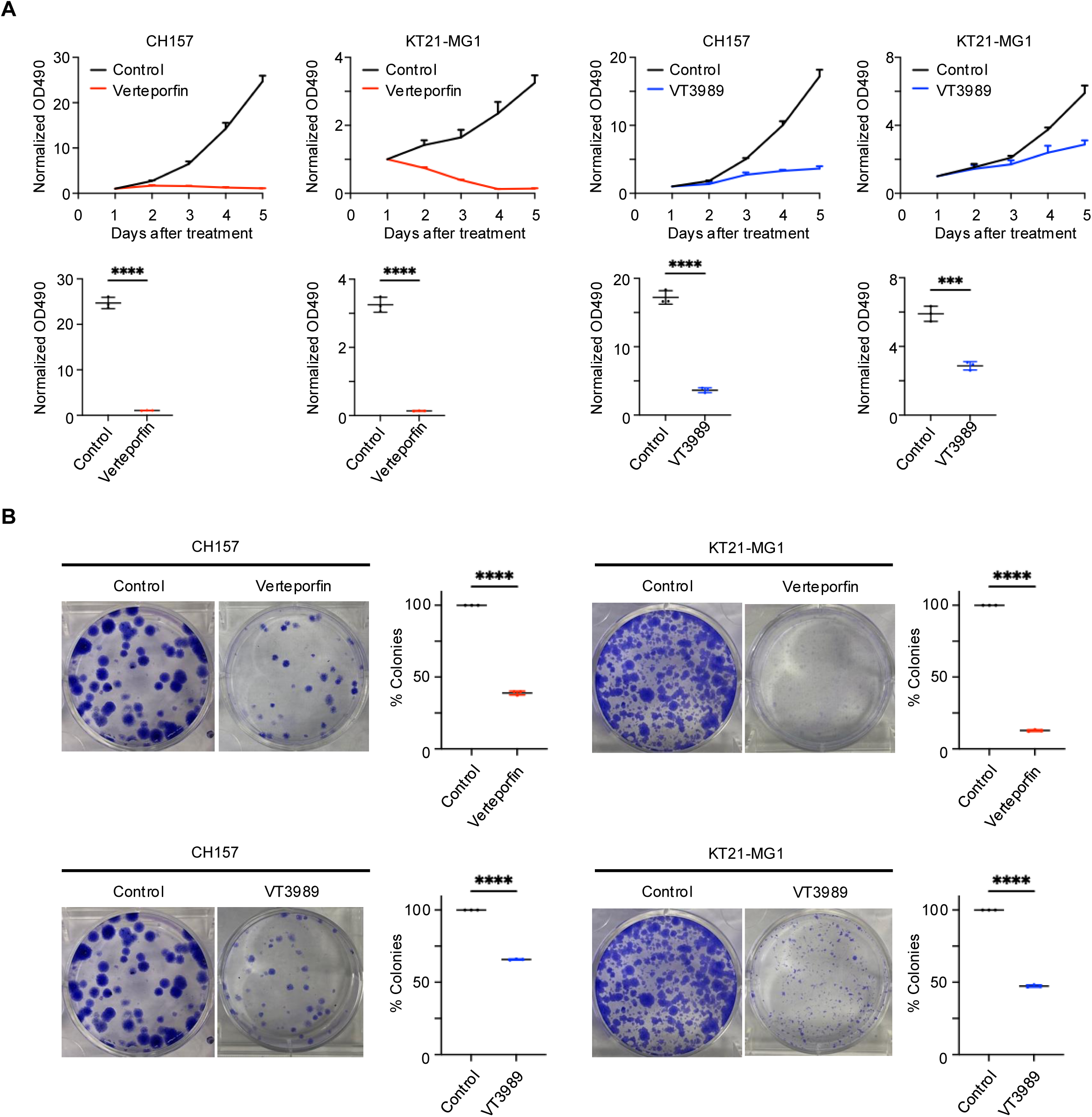
YAP–TEAD inhibition suppresses proliferation and clonogenic growth in CPA4-high meningioma cells. (A) Growth curves and day-5 OD490 values during treatment with vehicle, verteporfin, or VT3989 at the corresponding IC50 concentration. Data are mean ± SEM from three replicates normalized to day 1; unpaired, two-tailed t tests: verteporfin, CH157 *P* < 0.0001 and KT21-MG1 *P* < 0.0001; VT3989, CH157 *P* < 0.0001 and KT21-MG1 *P* = 0.0005. (B) Colony-formation assays and quantification. Data are mean ± SEM from three replicates; unpaired, two-tailed t tests, *P* < 0.0001 for each comparison.

**Supplementary Figure S5.**
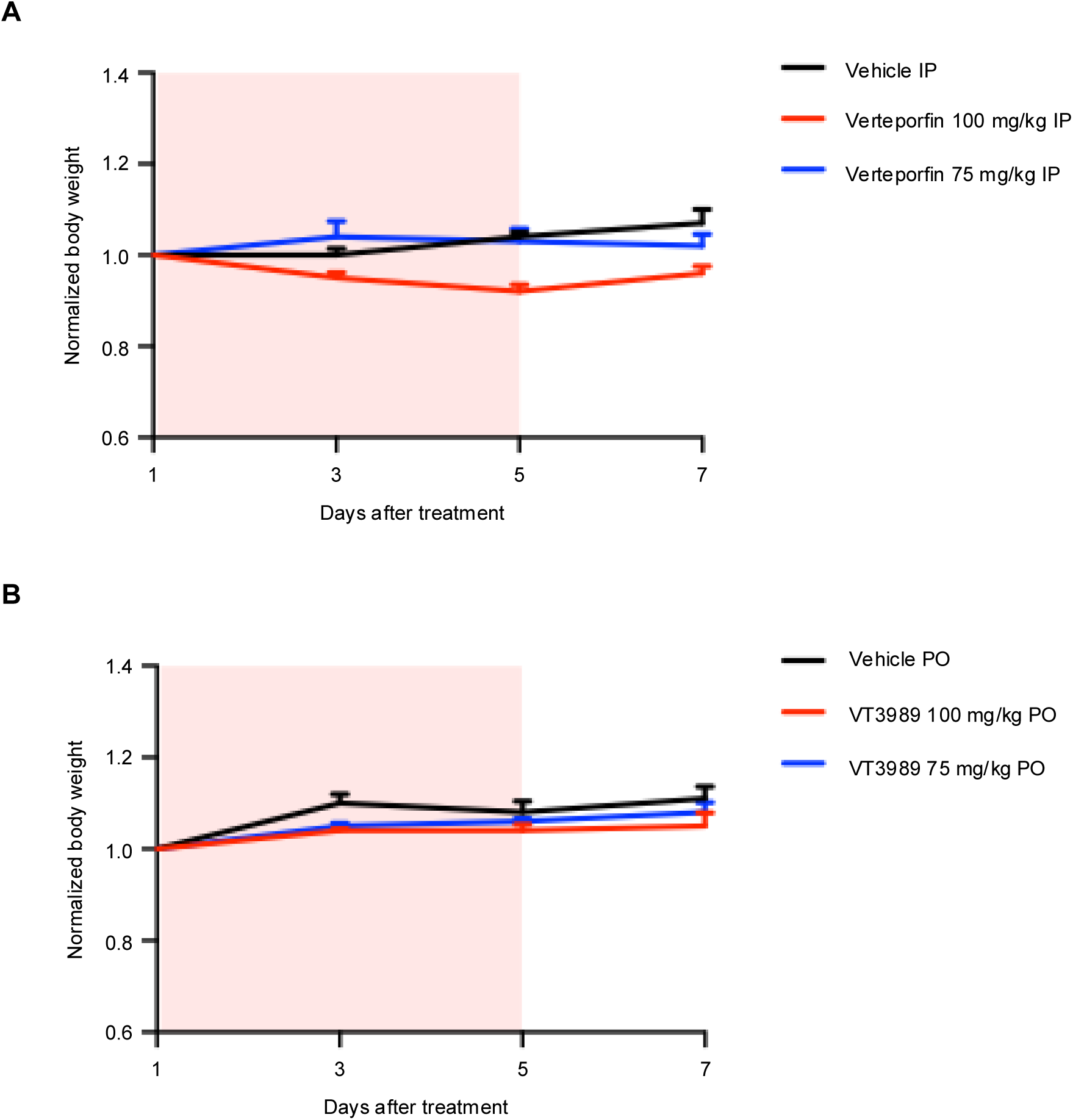
Dose tolerability of verteporfin and VT3989. (A) Body weight during treatment with vehicle or verteporfin (75 or 100 mg/kg intraperitoneally; n = 3 per group). (B) Body weight during treatment with vehicle or VT3989 (75 or 100 mg/kg orally; n = 3 per group). Agents were administered 5 days per week; shading denotes the treatment period.

**Supplementary Table S1.** Genes most strongly downregulated by NF2 restoration. The ten most strongly downregulated genes among those commonly downregulated in CH157 and KT21-MG1 cells are ranked by log2 fold change. Adjusted *P* values were calculated with DESeq2.

| Rank | KT21-MG1 |  |  | CH157 |  |  |
| --- | --- | --- | --- | --- | --- | --- |
|  | Gene | Log2FC | Padj | Gene | Log2FC | Padj |
| 1 | L1CAM | -3.01413 | 1.88E-25 | NLGN4X | -12.0313 | 9.64E-22 |
| 2 | THBS1 | -2.51564 | 1.75E-106 | PIK3C2B | -3.24585 | 6.04E-10 |
| 3 | VCAN | -2.4011 | 8.60E-14 | SLC16A9 | -3.04368 | 6.26E-34 |
| 4 | SEMA7A | -2.39376 | 5.87E-22 | ADORA2A | -2.82595 | 0.01404 |
| 5 | SERPINE1 | -2.38354 | 4.36E-17 | MBOAT2 | -2.64158 | 2.25E-181 |
| 6 | CPA4 | -2.36052 | 2.69E-113 | IGSF9B | -1.6874 | 0.0284894 |
| 7 | C6orf223 | -2.35415 | 3.49E-06 | FABP4 | -1.52747 | 5.44E-08 |
| 8 | LRP1 | -2.2618 | 1.28E-67 | SALL2 | -1.50346 | 6.12E-07 |
| 9 | ITPRIPL1 | -2.11944 | 0.0015074 | CPA4 | -1.36536 | 6.80E-30 |
| 10 | COL1A2 | -2.11393 | 1.01E-131 | C6orf223 | -1.34456 | 2.70E-06 |

**Supplementary Table S2.** TEAD motifs at the CPA4 promoter. Known TEAD motifs identified by HOMER analysis at the CPA4 promoter are shown with their corresponding sequences.

| Motif | Sequence |
| --- | --- |
| TEAD1(TEAD)/HepG2-TEAD1-ChIP-Seq | ATAAATTCCA |
| TEAD3(TEA)/HepG2-TEAD3-ChIP-Seq | TAAATTCCAT |
| TEAD(TEA)/Fibroblast-PU.1-ChIP-Seq | AAATTCCATC |

